# Whole genome genetic variation and linkage disequilibrium in a diverse collection of *Listeria monocytogenes* isolates

**DOI:** 10.1101/2020.11.02.364679

**Authors:** Swarnali Louha, Richard J. Meinersmann, Travis C. Glenn

**Author notes:** Corresponding author (SL).

## Abstract

We performed whole-genome multi-locus sequence typing for 2554 genes in a large and heterogenous panel of 180 *Listeria monocytogenes* strains having diverse geographical and temporal origins. The subtyping data was used for characterizing genetic variation and evaluating patterns of linkage disequilibrium in the pan-genome of *L. monocytogenes*. Our analysis revealed the presence of strong linkage disequilibrium in *L. monocytogenes*, with ∼99% of genes showing significant non-random associations with a large majority of other genes in the genome. Twenty-seven loci having lower levels of association with other genes were considered to be potential “hot spots” for horizontal gene transfer (i.e., recombination via conjugation, transduction, and/or transformation). The patterns of linkage disequilibrium in *L. monocytogenes* suggest limited exchange of foreign genetic material in the genome and can be used as a tool for identifying new recombinant strains. This can help understand processes contributing to the diversification and evolution of this pathogenic bacteria, thereby facilitating development of effective control measures.

## Introduction

The bacterial genome is a dynamic structure. Characterizing patterns of genomic variation in bacterial pathogens can provide insights into the forces shaping their biology and evolutionary history (Zwick et al 2011). Homologous recombination is an important driver of evolution and increases the adaptive potential of bacteria by allowing variation to be tested across multiple genomic backgrounds (Yahara et al. 2015). Recombination is mediated by three mechanisms; transformation, transduction, and conjugation, and the availability and efficacy of these mechanisms and their biological consequences play a major role in determining the frequency of recombination in a bacterial population (Feil and Spratt 2001, Zwick et al 2011). Recombination is variably distributed in bacterial genomes, with some sites in the genome recombining at a higher or lower frequency than the genomic average, known as hot spots and cold spots respectively (Steiner and Smith 2005). Evidence for recombination and its effect on genomic variation can be obtained by detecting patterns of non-random association of genotypes at different loci within a given population, termed as linkage disequilibrium (Feil and Spratt 2001, Zwick et al 2011). Various methods for detecting linkage disequilibrium have been used to study the extent of genetic recombination shaping the population structures of several bacterial species (Smith et al. 1993, Zwick et al. 2011, Takuno et al. 2012, Vigué and Eyre-Walker 2019).

*Listeria monocytogenes*, known for causing life-threatening infections in animals and human populations at risk, is one of the bacterial species having the lowest rate of homologous recombination. Genetic diversity in this species is mainly driven by the accumulation of mutations over time, with alleles five times more likely to change by mutation than by recombination (Ragon et al. 2008). *L. monocytogenes* is generally considered to have a clonal genetic structure (Piffaretti et al. 1989, Wiedmannn et al. 1997). The population structure of this bacteria consists of 4 evolutionary lineages (I, II, III and IV) and recombination has been observed between isolates of different lineages; suggesting that although recombination is rare in *L. monocytogenes*, this species is not completely clonal (den Bakker et al. 2008, Dunn et al. 2009, Ragon et al. 2008). Interestingly, homologous recombination is not equally frequent among isolates of different lineages, with lineages II, III and IV showing higher rates of recombination and lower degree of sequence similarity than lineage I (Meinersmann et al. 2004, den Bakker et al. 2008, Orsi et al. 2008, Kuenne et al. 2013).

Whole-genome sequencing studies have shown that *L. monocytogenes* genomes are highly syntenic in their gene content and organization, with a majority of gene-scale differences occurring in the accessory genome and accumulated in a few hypervariable hotspots, prophages, transposons, scattered unique genes and genetic islands encoding proteins of unknown functions (Nelson et al. 2004, Hain et al. 2007, den Bakker et al. 2010, 2013, Kuenne et al. 2013). Several other studies have detected evidence of recombination using a few genes (den Bakker et al. 2008, Cantinelli et al. 2013, Ragon et al. 2008) and indicated the presence of significant linkage disequilibrium in *L. monocytogenes* (Call et al. 2003, Salcedo et al. 2003). However, these studies used a limited number of *L. monocytogenes* isolates and evaluated recombination present in a small fraction of the genome, mostly made up of house-keeping genes, which are assumed to be under negative selection and less subject to homologous recombination.

Prior to the advent of next-generation sequencing technologies, multi locus enzyme electrophoresis (MLEE), was used for generating large data sets for the statistical analysis of bacterial populations. MLEE differentiates organisms by assessing the relative electrophoretic mobilities of intracellular enzymes and indexes allelic variation in multiple chromosomal genes (Mallik 2014). MLEE has been successfully used for studying the extent of linkage disequilibrium in a variety of bacterial species (Piffaretti et al. 1989, Maynard Smith et al. 1993, O’Rourke and Stevens 1993). With the easy and cheap availability of sequencing data in the last decade, MLEE has been replaced with an analogous technique called MLST (multi locus sequence typing) for subtyping bacterial genomes (Salcedo et al. 2003, Moura et al. 2017). We recently provided an approach that can generate whole-genome MLST (wgMLST) based characterization of *L. monocytogenes* isolates from whole-genome sequencing data (Louha et al. 2020). In this study, we use this wgMLST-based approach for characterizing genomic variation and assessing genome-wide patterns of linkage disequilibrium in a large collection of *L. monocytogenes* isolates obtained from diverse ecological niches.

## Materials and methods

### L. monocytogenes isolate selection

We selected a large and diverse panel of 180 *L. monocytogenes* isolates collected from different ecological communities (File S1). This set included (i) 20 isolates each from food, food contact surfaces (FCS), manure, milk, clinical cases, soil, and ready-to-eat (RTE) products obtained from the NCBI Pathogen Detection database and, (ii) 20 isolates from water and sediment samples in the South Fork Broad River watershed located in Northeast Georgia and 20 isolates from effluents from poultry processing plants (EFPP), provided by the USDA and FSIS.

### Whole-genome multi-locus sequence typing (wgMLST)

Whole-genome sequencing data for the 180 *L. monocytogenes* isolates were processed using Haplo-ST (Louha et al. 2020) for allelic profiling of 2554 genes per isolate. Illumina whole-genome sequencing reads obtained as previously described (File S1) were trimmed to remove all bases with a Phred quality score of < 20 from both ends and filtered such that 90% of bases in the clean reads had a quality of at least 20. After trimming and filtering, all remaining reads with lengths of < 50 bp were filtered out. The cleaned reads were assembled into allele sequences with YASRA (Ratan 2009) by mapping to reference genes and provided wgMLST profiles with BIGSdb-*Lm* (available at http://bigsdb.pasteur.fr/listeria).

### Analysis of Linkage Disequilibrium

First, the raw wgMLST profiles were filtered to remove paralogous loci and genes were ordered according to their genomic position in the *L. monocytogenes* reference strain EGD-e (NCBI accession number NC_003210.1). Next, new alleles not defined in the BIGSdb-*Lm* database and reported as ‘closest matches’ to existing alleles in BIGSdb-*Lm* were assigned custom allele ID’s with in-house Python scripts. The wgMLST profiles were further filtered to retain loci with < 5% missing data. The remaining loci were used to evaluate linkage disequilibrium (LD) between all pairs of loci with Arlequin v3.5.2 (Excoffier and Lischer 2010). LD tests for the presence of significant statistical association between pairs of loci and is based on an exact test. The test procedure is analogous to Fisher’s exact test on a two-by-two contingency table but extended to a contingency table of arbitrary size (Slatkin 1994). For each pair of loci, first a contingency table is constructed. The *k*_*1*_ x *k*_*2*_ entries of this table are the observed haplotype frequencies, with *k*_*1*_ and *k*_*2*_ being the number of alleles at locus 1 and locus 2, respectively. The LD test consists in obtaining the probability of finding a table with the same marginal totals and which has a probability equal or less than that of the observed contingency table. Instead of enumerating all possible contingency tables, a Markov chain is used to explore the space of all possible tables. To start from a random initial position in the Markov chain, the chain is explored for a pre-defined number of steps (the dememorization phase), such as to allow the Markov chain to forget its initial phase and make it independent from its starting point. The *P*-value of the test is then taken as the proportion of the visited tables having a probability smaller or equal to the observed contingency table. In our analysis, we used 100,000 steps of Markov chain to test the *P*-value of the LD test and 10,000 dememorization steps to reach a random initial position on the Markov chain. The significance level of the LD test was set at a *P*-value of 0.05.

### Assessment of genetic diversity

Genetic diversity between *L. monocytogenes* isolates collected from the different ecological niches listed as the isolate sources (File S1) was computed with pairwise F_ST_’s in Arlequin. F_ST_ measures the proportion of the variance in allele frequencies attributable to variation between populations (Charlesworth and Charlesworth 2010) and has a history of being used as a measure of the level of differentiation between populations in population genetics. Fifty thousand permutations were used to test the significance of the genetic distances at a significance level of 0.05.

The AMOVA procedure in Arlequin was used to compute the pairwise differences in allelic content between isolate wgMLST profiles as a matrix of Euclidean squared distances. This distance matrix was used to compute a minimum spanning tree (MST) between all isolates. The MST was visualized and annotated with iTOL v3 (Letunic and Bork 2016). For better visualization, the MST was converted to circular format and annotations for the source of isolates were displayed in outer external rings.

## Results

We performed whole-genome multi locus sequence typing for 180 *L. monocytogenes* isolates obtained from 9 different source populations. For each isolate, allele sequences were assembled for 2554 genes and provided allele ID’s based on the unified nomenclature available in the BIGSdb-*Lm* database (File S2). This dataset was filtered to remove 133 paralogous loci identified by Haplo-ST and all loci with > 5% missing data (alleles not assigned ID’s by Haplo-ST), and the remaining 2233 loci (File S3) were ordered according to their position in the *L. monocytogenes* reference genome EGD-e. Figure 1 shows the minimum spanning tree of the 180 isolates inferred from allelic differences in the wgMLST profiles. Two results are apparent. First, we see a long branch (red) containing a majority of isolates obtained from soil and manure clustered together, which suggests the origin of these strains from a common ancestor. Interestingly, 3 clinical strains are also found in this cluster. Secondly, a large number of food-related isolates (∼51%, obtained from food, FCS, RTE products and EFPP) clustered together in a single branch of the tree (blue) with short branch-lengths to the tips, suggesting that these strains are closely related to each other. Although this is expected, it is interesting to find a few strains obtained from clinical cases, river water and milk in this cluster. The presence of isolates from unrelated ecological communities could be due to the technique used for constructing the dendrogram, which groups isolates based on pairwise differences in allelic content between isolate wgMLST profiles rather than characterizing differences between all variants in nucleotide sequences.

**Figure 1:**
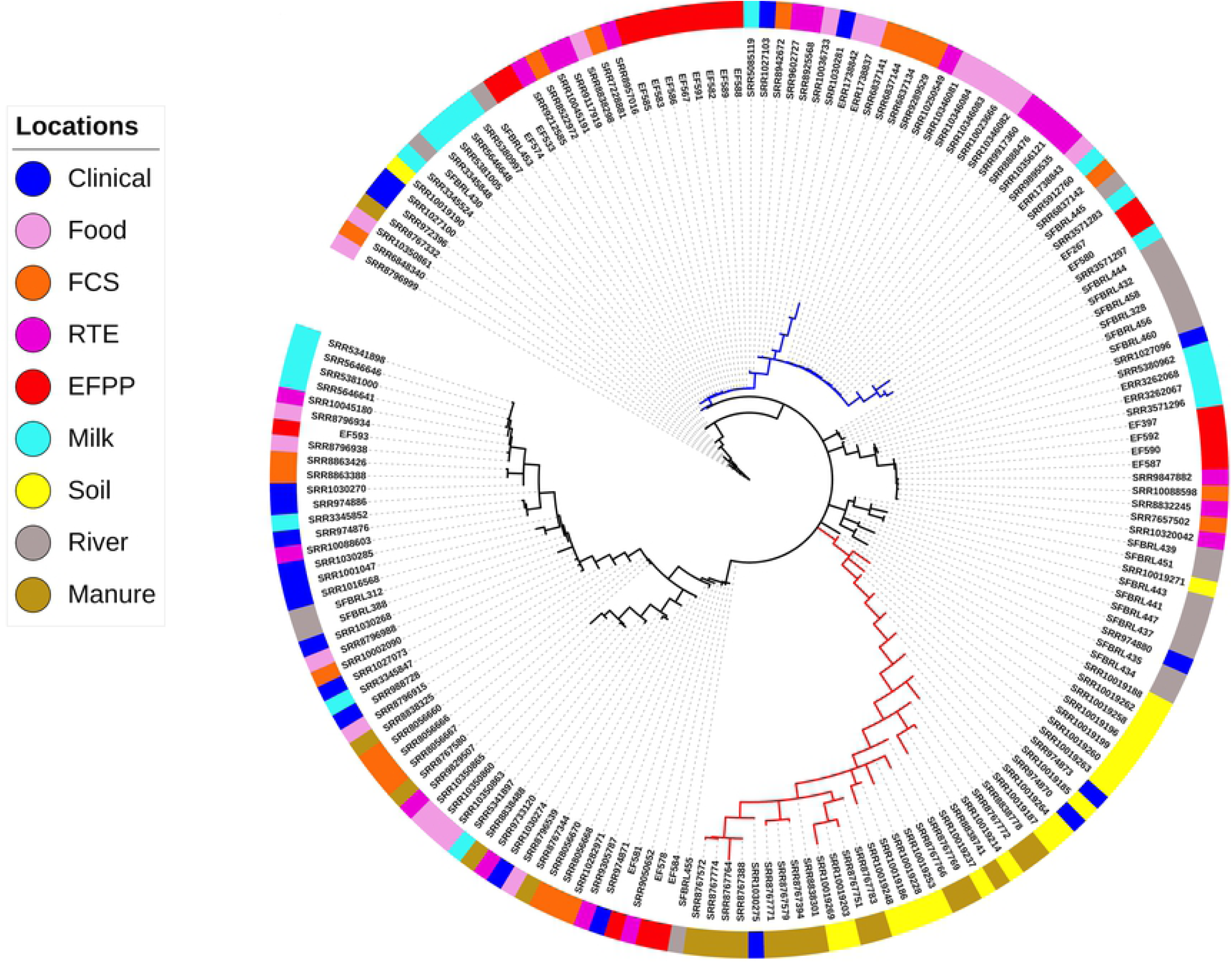
Patterns of genetic differentiation in the 180 *L. monocytogenes* isolates. Minimum spanning tree based on a distance matrix measuring pairwise differences in allelic content between isolate wgMLST profiles. The isolation source of each isolate is indicated with colors on the outer ring. Majority of the isolates sampled from soil and manure cluster together in a distant branch (red), suggesting their recent emergence from a common ancestor. A large number of food-related isolates cluster together in a single branch of the tree (blue), suggesting their close relatedness.

The genetic differentiation test that computes pairwise F_ST_’s between isolates collected from different ecological communities (Table 1) shows that isolates obtained from soil and manure show considerable genetic differentiation from isolates belonging to other communities, with the exception of isolates obtained from clinical cases. Secondly, isolates from the EFPP-RTE pairing has lower F_ST_ than EFPP pairing from all other locations. Thirdly, the clustering dendrogram (Fig 1) and F_ST_ test are supportive of each other in that isolates from RTE, FCS and food are not distinguished as separate populations.

**Table 1:**
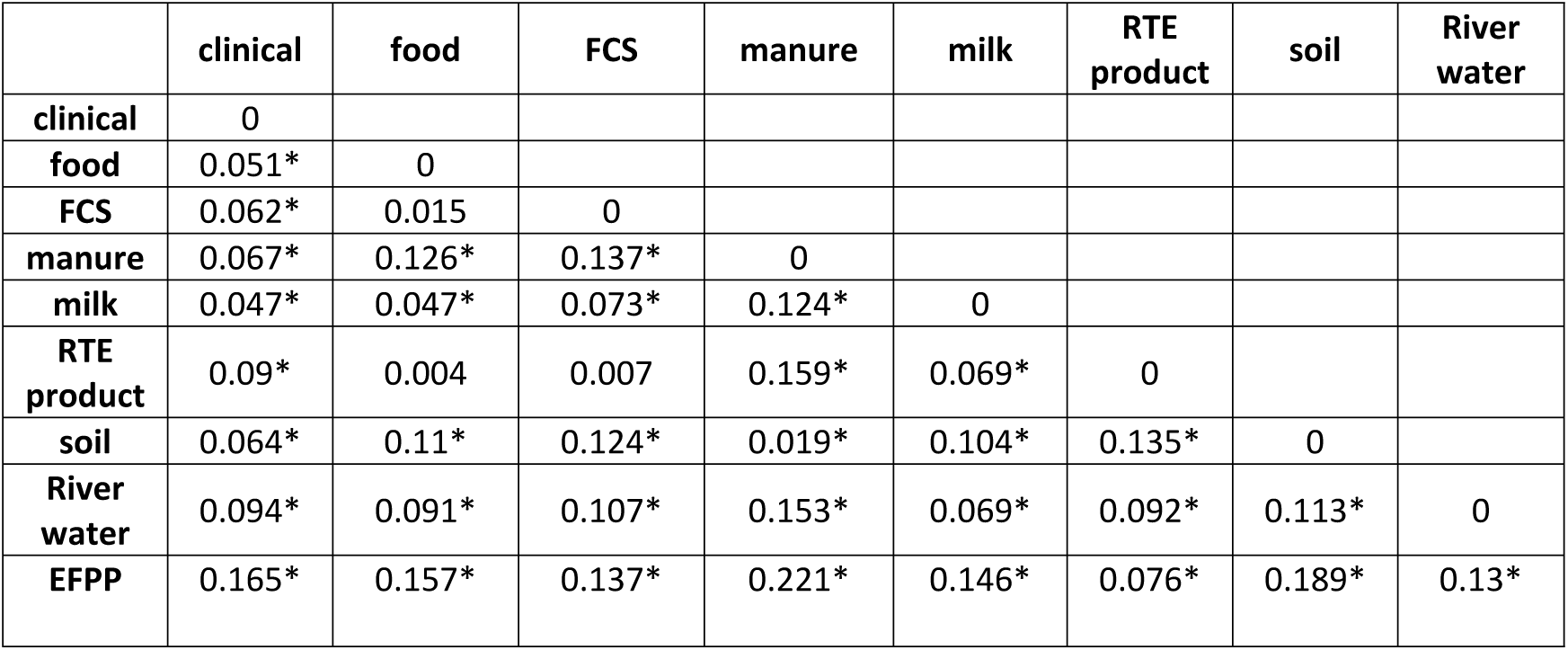
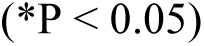
Pairwise genetic distances (F_ST_) between groups of *L. monocytogenes* strains isolated from nine different ecological niches.

|  | clinical | food | FCS | manure | milk | RTE product | soil | River water |
| --- | --- | --- | --- | --- | --- | --- | --- | --- |
| clinical | 0 |  |  |  |  |  |  |  |
| food | 0.051* | 0 |  |  |  |  |  |  |
| FCS | 0.062* | 0.015 | 0 |  |  |  |  |  |
| manure | 0.067* | 0.126* | 0.137* | 0 |  |  |  |  |
| milk | 0.047* | 0.047* | 0.073* | 0.124* | 0 |  |  |  |
| RTE product | 0.09* | 0.004 | 0.007 | 0.159* | 0.069* | 0 |  |  |
| soil | 0.064* | 0.11* | 0.124* | 0.019* | 0.104* | 0.135* | 0 |  |
| River water | 0.094* | 0.091* | 0.107* | 0.153* | 0.069* | 0.092* | 0.113* | 0 |
| EFPP | 0.165* | 0.157* | 0.137* | 0.221* | 0.146* | 0.076* | 0.189* | 0.13* |

(\*P < 0.05)

We investigated LD between pairs of genes in the genome using an exact test, which measures non-random associations between alleles at two loci based on the difference between observed and expected allele frequencies. As expected, most genes pairs (∼97%) in the genome of *L. monocytogenes* show significant LD among pairs of alleles (Fig 2, File S4). A majority of genes (2205 of of 2233, ∼99%) were found to be at LD with at least 90% of other genes in the genome (File S5). Of the remaining 27 genes (∼1%) that were at LD with < 90% of genes (Table 2), 10 genes were found to be at LD with < 50% of genes. A single locus, *lmo0046*, was at LD with only 19 other genes.

**Figure 2:**
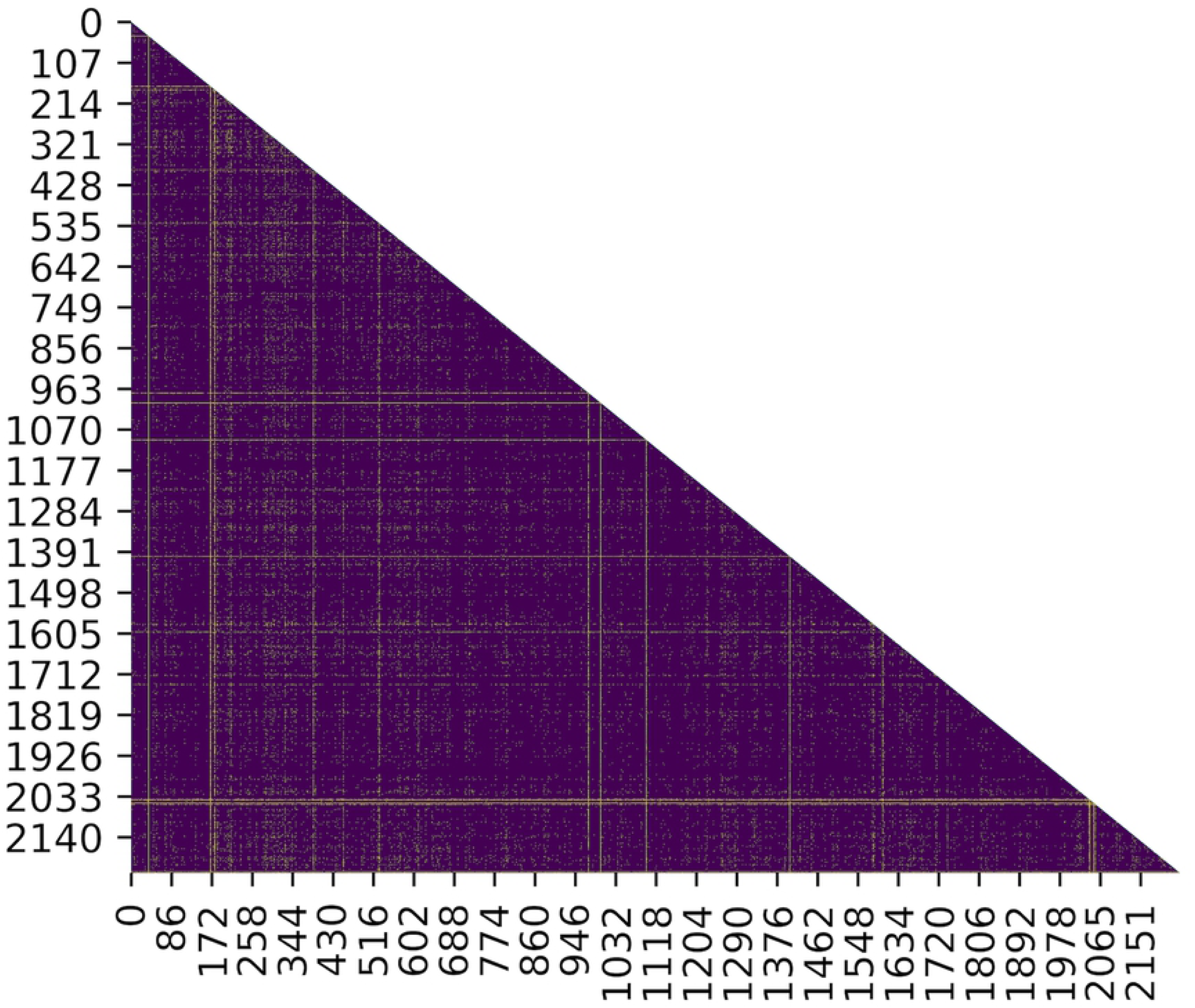
Heatmap of the extent of LD in the genome of *L. monocytogenes*. Genes are ordered according to their genomic positions in the L. monocytogenes reference strain EGD-e along the x and y axis (for gene names see File S4). A majority of genes show significant LD in the genome (indigo), while few genes are at linkage equilibrium (yellow).

**Table 2:** Genes at LD with < 90% of genes in the genome of *L. monocytogenes*, thus showing significant evidence for horizontal genetic transfer.

| Gene name | Number of genes at LD | Percentage of genes at LD | Location in the chromosome of EGD-e (bp) | Location in core/accessory genome* | Function |
| --- | --- | --- | --- | --- | --- |
| **lmo0046** | 19 | 0.85 | 50514..50753 | core | small subunit ribosomal protein S18 |
| **lmo2624** | 185 | 8.289 | 2701254..2701445 | core | large subunit ribosomal protein L29 |
| **lmo2856** | 215 | 9.63 | 2943569..2943703 | accessory | large subunit ribosomal protein L34 |
| **lmo1364** | 239 | 10.71 | 1387014..1387214 | accessory | Cold shock protein |
| **lmo1469** | 454 | 20.34 | 1501881..1502054 | core | small subunit ribosomal protein S21 |
| **lmo2616** | 458 | 20.52 | 2697988..2698347 | accessory | large subunit ribosomal protein L18 |
| **lmo1816** | 484 | 21.69 | 1890951..1891139 | core | large subunit ribosomal protein L28 |
| **lmo0248** | 576 | 25.81 | 265029..265454 | accessory | large subunit ribosomal protein L11 |
| **lmo1335** | 880 | 39.43 | 1363826..1363975 | core | large subunit ribosomal protein L33 |
| **inlH (lmo0263)** | 1006 | 45.07 | 284365..286011 | accessory | internalin H |
| **cwhA (lmo0582)** | 1223 | 54.79 | 618932..620380 | accessory | Invasion associated secreted endopeptidase |
| **lmo2047** | 1377 | 61.69 | 2130228..2130401 | accessory | large subunit ribosomal protein L32 |
| **lmo2628** | 1508 | 67.56 | 2702909..2703187 | accessory | small subunit ribosomal protein S19 |
| **lmo2614** | 1580 | 70.79 | 2697267..2697446 | core | large subunit ribosomal protein L30 |
| **lmo0758** | 1606 | 71.95 | 783901..784788 | core | Hypothetical protein |
| **lmo0514** | 1699 | 76.12 | 547520..549337 | accessory | Internalin |
| **lmo0305** | 1709 | 76.57 | 329923..330999 | core | L-allo-threonine aldolase |
| **lmo0659** | 1771 | 79.35 | 699410..700306 | accessory | Transcriptional regulator |
| **lmo2206** | 1791 | 80.24 | 2294555..2297155 | accessory | Heat shock proteins |
| **Imo0756** | 1797 | 80.51 | 781896..782801 | core | ABC Transporters |
| **Imo0865** | 1859 | 83.29 | 903837..905510 | core | Amino sugar and nucleotide sugar metabolism |
| **Imo2014** | 1888 | 84.59 | 2088797..2091454 | accessory | Glycan biosynthesis and metabolism |
| **Imo1611** | 1904 | 85.3 | 1654902..1655975 | core | Aminopeptidase |
| **inIE (Imo0264)** | 1913 | 85.71 | 286219..287718 | accessory | Internalin E |
| **Imo1839** | 1925 | 86.25 | 1916166..1917452 | accessory | Electrochemical potential-driven transporters |
| **Imo2179** | 1968 | 88.17 | 2264772..2268230 | accessory | Peptidoglycan binding protein |
| **inIB (Imo0434)** | 1981 | 88.75 | 457021..458913 | accessory | Internalin B |

\*Location in core/accessory genome has been determined with respect to the core-genome
MLST scheme developed by the Institut Pasteur (Moura et al. 2016 Nat Microbiol 2: 16185).

## Discussion

Our dataset reveals the presence of strong LD in the genome of *L. monocytogenes*. Among the 2233 genes tested for LD, 2205 genes (approx. 99%) were found to have pairwise LD with a majority of other genes (90%) in the genome. High levels of LD can not only arise in highly clonal bacterial populations with low rates of recombination, but may also be temporarily present in bacteria with ‘epidemic’ population structures, in which high recombination rates randomize association between alleles, but adaptive clones emerge and diversify over the short-term (Smith et al. 1993, Feil and Spratt 2001). Because *Listeria* has a clonal genetic structure, it is difficult to see how this high level of LD can arise except as a consequence of low rates of recombination. This is consistent with studies which report recombination in chromosomal genes as an infrequent event in natural populations of *L. monocytogenes* (Piffaretti et al. 1989, Ragon et al. 2008). Because the extent of genetic linkage is a useful index to the horizontal transfer occurring within a species and can be presented as direct evidence for recombination (Feil et al. 2001), the remaining ∼1% of genes (Table 2) that were at LD with < 90% of genes can be described as “hot spots” for the gain of horizontally acquired information. The extensive linkage disequilibrium that we describe in *L. monocytogenes* is in sharp contrast to other pathogenic bacteria that are naturally competent for transformation and recombine frequently to give rise to either weakly clonal or panmictic population structures (Duncan et al. 1994, Suerbaum et al. 1998, Al Suwayyid et al. 2018).

The *L. monocytogenes* pan-genome is highly conserved but open to limited acquisition of foreign DNA or genetic variability through evolutionary forces such as mutation, duplication or recombination (Kuenne et al. 2013). Evidence for homologous recombination between closely related strains of *L. monocytogenes* has been detected by multiple studies, however, non-homologous recombination seems to be rare (Orsi et al. 2008, Dunn et al. 2009, Nightingale et al. 2005). Although recombination via conjugation and generalized transduction has been reported in *L. monocytogenes* (Flamm et al. 1984, Lebrun et al. 1992, Hodgson 2000), and most competence related genes are present in all *Listeria* genomes (Buchrieser 2007), natural competence or induced competence under laboratory conditions has not been observed in *L. monocytogenes* (Borezee et al. 2000, Glaser et al. 2001). This lack of competence may partially explain the low levels of gene acquisition from external gene pools. Limited gene acquisition may also be facilitated by defense systems for foreign DNA/mobile elements such as restriction-modification and/or CRISPR systems, both of which have been shown to restrict horizontal gene transfer in other bacterial genera (den Bakker et al. 2010).

The frequency of recombination in *L. monocytogenes* differs considerably in different regions of the genome and between isolates of different lineages (den Bakker et al. 2008, 2013). This may arise from differences in selective pressures in the environment and varying degrees of horizontal gene transfer. Several comparative genomic studies report a clustered distribution of accessory genes on the right replichore of the *L. monocytogenes* genome (approx. 500 Kb in the first 65°), indicating an area of high genome plasticity (Kuenne et al. 2013, den Bakker et al. 2013). On the contrary, a study by Orsi et al. failed to find any evidence of spatial clustering in a large number of genes which show evidence for recombination in *L. monocytogenes* (Orsi et al. 2008). Further, a recent study described the presence of homologous recombination in nearly 60% of loci in the core genome of *L. monocytogenes*, although most of this variation was also found to be affected by purifying selection and was thus neutral (Moura et al. 2016). This is consistent with results from our analysis which finds linkage equilibrium between only ∼1% of gene pairs in the genome. Also, genes considered as potential recombination hot spots (Table 2) in our dataset are found to be scattered in the genome. A large number (∼41%) of these “hot spot” genes (*lmo0046, lmo2624, lmo2856, lmo1469, lmo2616, lmo1816, lmo0248, lmo1335, lmo2047, lmo2628, lmo2614*), encode ribosomal proteins and their related subunits. According to the complexity theory (Jain et al. 1999), informational genes involved in complex biosystems and maintenance of basal cellular functions are usually conserved, as they might be less likely to be compatible in the systems of other species. Thus, housekeeping genes such as ribosomal proteins are generally considered to be relatively restricted to horizontal gene transfer. However, several reports suggest horizontal gene transfer of ribosomal proteins in many prokaryotic genomes (Brochier et al. 2000, Makarova et al. 2001, Garcia-Vallve et al. 2002, Chen et al. 2009). Two other “hot spot” genes (*lmo0865, lmo2014*) are involved in carbohydrate and amino acid metabolism and have shown evidence for recombination in a prior study (Orsi et al. 2008), indicating that the rapid diversification of these genes may enable *L. monocytogenes* to adapt to environments with varying nutrient availabilities. Some of the other genes encode a variety of internalin’s (*lmo0263, lmo0514, lmo0264*, lmo0434), transporters (*lmo0756, lmo1839*), transcriptional regulators (*lmo0659*), cell surface proteins (*lmo2179*), other invasion-associated proteins (*lmo0582*), and proteins involved in response to temperature fluctuations (*lmo1364, lmo2206*). Internalin’s are cell surface proteins with known and hypothesized roles in virulence (den Bakker et al. 2010, Tsai et al. 2011). Evidence of recombination in internalin’s and these other genes suggests that *L. monocytogenes* is subjected to sustained selection pleasures in the environment, and it responds to these pressures by continuously regulating its transcriptional machinery and remodeling the cell surface, thereby facilitating adaptation within the host and as a saprophyte.

In conclusion, we have identified the presence of strong linkage disequilibrium in the genome of *L. monocytogenes*. Parts of the genome showing strong non-random association between genes are highly conserved regions, and are most possibly affected by positive selection. The low levels of recombination within the *L. monocytogenes* genome suggests that the patterns of association observed between genes can be used to recognize newly emerging strains and help in understanding the processes involved in the diversification and evolution of *L. monocytogenes*. Such investigations can ultimately help to develop better control measures for this pathogenic microbe.

## Acknowledgments

We thank USDA and FSIS for providing us with *Listeria monocytogenes* whole-genome sequencing samples from river water and effluents of poultry processing plants. The high-performance computing cluster at Georgia Advanced Computing Resource Center (GACRC) at the University of Georgia provided computational infrastructure and technical support throughout the work.

## Supporting information

**S1 File**. Panel of 180 *L. monocytogenes* isolates collected from different ecological communities.

**S2 File**. Whole-genome MLST profiles of the 180 *L. monocytogenes* isolates.

**S3 File**. Whole-genome MLST profiles of 2233 loci retained for AMOVA after filtering out paralogous loci and loci with > 5% of missing data.

**S4 File**. Heatmap of LD in the genome of *L. monocytogenes*.

**S5 File**. Percentage of genes at LD with each gene in the genome of *L. monocytogenes*.

## References

1) Zwick ME, Thomason MK, Chen PE, Johnson HR, Sozhamannan S, Mateczun A et al. Genetic variation and linkage disequilibrium in Bacillus anthracis. Sci Rep. 2011;1: 169.

2) Yahara K, Didelot X, Jolley KA, Kobayashi I, Maiden MC, Sheppard SK et al. The Landscape of Realized Homologous Recombination in Pathogenic Bacteria. Mol Biol Evol. 2016;33: 456–471.

3) Feil EJ, Spratt BG. Recombination and the population structures of bacterial pathogens. Annu Rev Microbiol. 2001;55: 561–590.

4) Steiner WW, Smith GR. Natural meiotic recombination hot spots in the Schizosaccharomyces pombe genome successfully predicted from the simple sequence motif M26. Mol Cell Biol. 2005;25: 9054–9062.

5) Smith JM, Smith NH, O’Rourke M, Spratt BG. How clonal are bacteria? Proc Natl Acad Sci USA. 1993;90: 4384–4388.

6) Takuno S, Kado T, Sugino RP, Nakhleh L, Innan H. Population Genomics in Bacteria: A Case Study of Staphylococcus aureus. Mol Biol Evol. 2012;29: 797–809.

7) Vigué L, Eyre-Walker A. The comparative population genetics of Neisseria meningitidis and Neisseria gonorrhoeae. PeerJ. 2019;7: e7216. doi: 10.7717/peerj.7216.

8) Piffaretti JC, Kressebuch H, Aeschbacher M, Bille J, Bannerman E, Musser JM et al. Genetic characterization of clones of the bacterium Listeria monocytogenes causing epidemic disease. Proc Natl Acad Sci USA. 1989;86: 3818–3822.

9) Mallik S. IDENTIFICATION METHODS | Multilocus Enzyme Electrophoresis. In: Batt CA, Tortorello ML, editors. Reference Module in Food Science: Encyclopedia of Food Microbiology (Second Edition); 2014. Pp. 336–343.

10) O’Rourke M, Stevens E. Genetic structure of Neisseria gonorrhoeae populations : a nonclonal pathogen. Journal of General Microbiology. 1993;139: 2603–2611.

11) Moura A, Criscuolo A, Pouseele H, Maury MM, Leclercq A, Tarr C et al. Whole genome-based population biology and epidemiological surveillance of Listeria monocytogenes. Nat Microbiol. 2016;2: 16185.

12) Wiedmann M, Bruce JL, Keating C, Johnson AE, McDonough PL, Batt CA. Ribotypes and Virulence Gene Polymorphisms Suggest Three Distinct Listeria monocytogenes Lineages With Differences in Pathogenic Potential. Infect Immun. 1997;65: 2707–2716.

13) Call DR, Borucki MK, Besser TE. Mixed-genome Microarrays Reveal Multiple Serotype and Lineage-Specific Differences Among Strains of Listeria monocytogenes. J Clin Microbiol. 2003;41: 632–639.

14) Ragon M, Wirth T, Hollandt F, Lavenir R, Lecuit M, Le Monnier A et al. A new perspective on Listeria monocytogenes evolution. PLoS Pathog. 2008;4(9): e1000146. doi: 10.1371/journal.ppat.1000146.

15) den Bakker HC, Didelot X, Fortes ED, Nightingale KK, Wiedmann M. Lineage Specific Recombination Rates and Microevolution in Listeria monocytogenes. BMC Evol Biol. 2008;8: 277.

16) Dunn KA, Bielawski JP, Ward TJ, Urquhart C, Gu H. Reconciling Ecological and Genomic Divergence Among Lineages of Listeria Under an “Extended Mosaic Genome Concept”. Mol Biol Evol. 2009;26: 2605–2615.

17) Orsi RH, Sun Q, Wiedmann M. Genome-wide analyses reveal lineage specific contributions of positive selection and recombination to the evolution of Listeria monocytogenes. BMC Evol Biol. 2008;8: 233.

18) Kuenne C, Billion A, Mraheil MA, Strittmatter A, Daniel R, Goesmann A et al. Reassessment of the Listeria monocytogenes Pan-Genome Reveals Dynamic Integration Hotspots and Mobile Genetic Elements as Major Components of the Accessory Genome. BMC Genomics. 2013; 14:47.

19) Nelson KE, Fouts DE, Mongodin EF, Ravel J, DeBoy RT, Kolonay JF et al. Whole Genome Comparisons of Serotype 4b and 1/2a Strains of the Food-Borne Pathogen Listeria monocytogenes Reveal New Insights Into the Core Genome Components of This Species. Nucleic Acids Res. 2004;32: 2386–2395.

20) Hain T, Chatterjee SS, Ghai R, Kuenne CT, Billion A, Steinweg C et al. Pathogenomics of Listeria spp. Int J Med Microbiol. 2007;297: 541–557.

21) den Bakker HC, Cummings CA, Ferreira V, Vatta P, Orsi RH, Degoricija L et al. Comparative genomics of the bacterial genus Listeria: Genome evolution is characterized by limited gene acquisition and limited gene loss. BMC Genomics. 2010;11: 688.

22) den Bakker HC, Desjardins CA, Griggs AD, Peters JE, Zeng Q, Young SK et al. Evolutionary Dynamics of the Accessory Genome of Listeria monocytogenes. PLoS One. 2013;8: e67511. doi: 10.1371/journal.pone.0067511.

23) Cantinelli T, Chenal-Francisque V, Diancourt L, Frezal L, Leclercq A, Wirth T et al. “Epidemic clones” of Listeria monocytogenes are widespread and ancient clonal groups. J Clin Microbiol. 2013;51: 3770–3779.

24) Salcedo C, Arreaza L, Alcalá B, de la Fuente L, Vázquez JA. Development of a multilocus sequence typing method for the analysis of Listeria monocytogenes clones. J Clin Microbiol. 2003;41: 757–762.

25) Louha S, Meinersmann RJ, Abdo Z, Berrang ME, Glenn TC. An open-source program (Haplo-ST) for whole-genome sequence typing shows extensive diversity of Listeria monocytogenes in outdoor environments and poultry processing plants. Appl Environ Microbiol. 2020;87. DOI: 10.1128/AEM.02248-20

26) Excoffier L, Lischer HEL. Arlequin suite ver 3.5: A new series of programs to perform population genetics analyses under Linux and Windows. Mol Ecol Resour. 2010;10: 564–567.

27) Ratan A. Assembly algorithms for next generation sequence data. Ph.D. Dissertation, The Pennsylvania State University. 2009. Available from: https://etda.libraries.psu.edu/files/final_submissions/587

28) Letunic I, Bork P. Interactive tree of life (iTOL) v3: an online tool for the display and annotation of phylogenetic and other trees. Nucleic Acids Res. 2016; 44: W242–245.

29) Slatkin M. Linkage disequilibrium in growing and stable populations. Genetics. 1994;137: 331–336.

30) Charlesworth B, Charlesworth D. Elements of Evolutionary Genetics. Greenwood Village, Colorado: Roberts and Company publishers; 2010.

31) Duncan KE, Ferguson N, Kimura K, Zhou X, Istock CA. Fine-scale genetic and phenotypic structure in natural populations of Bacillus subtilis and Bacillus licheniformis: implications for bacterial evolution and speciation. Evolution. 1994;48: 2002–2025.

32) Al Suwayyid BA, Coombs GW, Speers DJ, Pearson J, Wise MJ, Kahler CM. Genomic epidemiology and population structure of Neisseria gonorrhoeae from remote highly endemic Western Australian populations. BMC Genomics. 2018;19: 165.

33) Suerbaum S, Smith JM, Bapumia K, Morelli G, Smith NH, Kunstmann E et al. Free recombination within Helicobacter pylori. Proc Natl Acad Sci USA. 1998;95: 12619–12624.

34) Buchrieser C. Biodiversity of the species Listeria monocytogenes and the genus Listeria. Microbes Infect. 2007;9: 1147–1155.

35) Nightingale K, Windham K, Wiedmann M. Evolution and molecular phylogeny of Listeria monocytogenes isolated from human and animal listeriosis cases and foods. J Bacteriol. 2005;187: 5537–5551.

36) Glaser P, Frangeul L, Buchrieser C, Rusniok C, Amend A, Baquero F et al. Comparative genomics of Listeria species. Science. 2001;294: 849–852.

37) Borezee E, Msadek T, Durant L, Berche P. Identification in Listeria monocytogenes of MecA, a homologue of the Bacillus subtilis competence regulatory protein. J Bacteriol. 2000;182: 5931–5934.

38) Jain R, Rivera MC, Lake JA. Horizontal gene transfer among genomes: the complexity hypothesis. Proc Natl Acad Sci USA. 1999;96: 3801–3806.

39) Garcia-Vallve S, Simo FX, Montero MA, Arola L, Romeu A. Simultaneous horizontal gene transfer of a gene coding for ribosomal protein l27 and operational genes in Arthrobacter sp. J Mol Evol. 2002;55: 632–637.

40) Makarova KS, Ponomarev VA, Koonin EV. Two C or not two C: recurrent disruption of Zn-ribbons, gene duplication, lineage-specific gene loss, and horizontal gene transfer in evolution of bacterial ribosomal proteins. Genome Biol. 2001;2: RESEARCH 0033. doi: 10.1186/gb-2001-2-9-research0033.

41) Chen K, Roberts E, Luthey-Schulten Z. Horizontal gene transfer of zinc and non-zinc forms of bacterial ribosomal protein S4. BMC Evol Biol. 2009;9: 179.

42) Brochier C, Philippe H, Moreira D. The evolutionary history of ribosomal protein RpS14: horizontal gene transfer at the heart of the ribosome. Trends Genet. 2000;16: 529–533.

43) Meinersmann RJ, Phillips RW, Wiedmann M, Berrang ME. Multilocus sequence typing of Listeria monocytogenes by use of hypervariable genes reveals clonal and recombination histories of three lineages. Appl Environ Microbiol. 2004;70: 2193–2203.

44) Flamm RK, Hinrichs DJ, Thomashow MF. Introduction of pAM beta 1 into Listeria monocytogenes by conjugation and homology between native L. monocytogenes plasmids. Infect Immun. 1984;44: 157–161.

45) Hodgson DA. Generalized transduction of serotype 1/2 and serotype 4b strains of Listeria monocytogenes. Mol. Microbiol. 2000;35: 312–323.

46) Lebrun M, Loulergue J, Chaslus-Dancla E, Audurier A. Plasmids in Listeria monocytogenes in relation to cadmium resistance. Appl Environ Microbiol. 1992;58: 3183–3186.

